# Functionality of the putative surface glycoproteins of the Wuhan spiny eel influenza virus

**DOI:** 10.1101/2021.01.04.425274

**Authors:** Guha Asthagiri Arunkumar, Shirin Strohmeier, Tiehai Li, Disha Bhavsar, Veronika Chromikova, Fatima Amanat, Mehman Bunyatov, Patrick C. Wilson, Ali H. Ellebedy, Geert-Jan Boons, Viviana Simon, Robert P. de Vries, Florian Krammer

## Abstract

A panel of novel influenza-like virus sequences were recently documented in jawless fish, ray-finned fish, and amphibians. Of these, the Wuhan spiny eel influenza virus (WSEIV) was found to phylogenetically cluster with influenza B viruses as a sister clade. Influenza B viruses have been historically documented to circulate only in humans, with certain virus isolates found in harbor seals. It is therefore interesting that a similar virus was potentially found in fish. Here we characterized the functionality and antigenicity of the putative hemagglutinin (HA) and neuraminidase (NA) surface glycoproteins of the WSEIV to better understand this virus and its pandemic potential. Upon functional characterization of NA, we identified that the WSEIV NA-like protein has sialidase activity comparable to B/Malaysia/2506/2004 influenza B virus NA, making it a *bona fide* neuraminidase that could be inhibited by NA inhibitors. Testing of the functionality of HA was carried out including receptor specificity, stability, and preferential airway protease cleavage and showed very specific binding to monosialic ganglioside 2 (GM2). To probe the degree of conservation of target epitopes, binding of known broadly cross-reactive monoclonal antibodies targeting the influenza B HA and NA, respectively, were assessed through enzyme linked immunosorbent assays against recombinant WSEIV glycoproteins. Human serum samples of patients with antibodies to influenza B viruses were used to determine the cross-reactivity against these novel glycoproteins. Very few monoclonal antibodies – notably including pan NA antibody 1G01 - showed cross-reactivity and reactivity from human sera was basically absent. In summary, we have conducted a functional and antigenic characterization of the glycoproteins of the novel WSEIV to assess if it is indeed a *bona fide* influenza virus potentially circulating in ray-finned fish.

## Introduction

Influenza A and B viruses cause widespread infections in humans on an annual basis resulting in significant morbidity and mortality.^1^ In addition to human infections, influenza A virus has been shown to have a broad host tropism, infecting a variety of different avian and mammalian species.^2^ This enhances the pandemic potential of these viruses due to an increased possibility of reassortment in a commonly infected host.^3^ In contrast, influenza B viruses have a comparatively limited tropism of hosts comprising predominantly of infections in humans. Sporadic outbreaks have been observed in harbor seals and gray seals, and this has brought into question the possibility of non-human reservoirs for influenza B viruses.^4,5^ However, sequence analysis of influenza B virus isolates from seals suggests that the causative agents were human strains.^6^ Partial sequences of isolates of influenza B viruses identified in swine farms across the USA displayed high homology to human influenza B virus isolates as well.^7^ Overall, studies have documented spillage of influenza B virus infections from humans to other species, however, there is limited evidence for the existence of sustained animal reservoirs for these viruses.

The overall understanding of RNA virus diversity outside of avian and mammalian species has been limited stemming from sampling biases towards these hosts.^8^ A study by Shi et al. aimed at addressing this dearth of sampling in amphibians, fish, and reptiles, and identified 214 vertebrate species-associated virus sequences through a meta-transcriptomic approach. Most of these viruses could be categorized into 17 vertebrate-specific viral families, significantly enhancing the diversity of viral families historically known to have mammalian or avian hosts. Three novel influenza viruses were identified in ray-finned fish (spiny eel), jawless fish (hagfish), and amphibians (Asiatic toad). These were the first documented sequences of putative influenza viruses in fish.^9^

For the Wuhan spiny eel influenza virus (WSEIV), all eight genomic segments were recovered following the sampling and analysis of the transcripts in the gill tissues of lesser spiny eels. A striking aspect of this virus is its phylogenetic clustering as a sister clade to influenza B viruses, more so than influenza A viruses do. Alignment of the coding regions of the eight segments of WSEIV indicates percentage identity as high as 76% (PB1) and as low as 34% (NS) with the closest hits all being in the influenza B virus family. The surface viral glycoproteins, hemagglutinin (HA) and the neuraminidase (NA) of the WSEIV, have a 45% and 48% amino acid identity to the respective HA and NA of influenza B viruses.^9^

Given our interest in studying viral glycoproteins, we decided to take a deeper dive into characterizing the HA and NA of the WSEIV and their influenza B virus counterparts. This could provide a valuable insight into this novel influenza B-like virus, and its implication of non-human reservoirs for influenza B viruses. Additionally, influenza B viruses are largely understudied relative to influenza A viruses in context of functionality and antigenic landscape, and the findings from this study contribute towards addressing this gap.^10^ The questions raised during the conception of this study were as follows. Does the WSEIV HA have sialic acid binding activity and what receptor specificity does it possess? Does the WSEIV neuraminidase have sialidase enzymatic activity? How do the functional and antigenic aspects of these proteins compare to those of influenza B virus HA and NAs and how does this inform the potential of this virus to spill over into humans?

Here, we show that the WSEIV HA and NA show opposing similarity profiles relative to an influenza B/Malaysia/2506/2004 virus HA and NA. The WSEIV HA appears to strongly interact with a unique gangliosidic receptor, displaying drastically different target receptor specificity compared to influenza B virus HAs. On the contrary, the WSEIV NA, showing sialidase activity, has a notably similar activity profile relative to the control influenza B virus NA. Additional functional characterization further reinforces this dichotomous nature of the HA and NA. Overall, we show that this WSEIV is indeed a *bona fide* influenza virus from a functional standpoint. We also address the antigenic epitope conservation on the WSEIV HA and NA using a panel of broadly cross-reactive monoclonal antibodies (mAb) and a set of serum samples from humans positive for influenza B virus.

## Results

Representative amino acid sequences of influenza A and B virus subtypes HA and NA proteins were selected and phylogenetically compared to the WSEIV HA and NA. Along the lines of the whole virus genome alignments in the study identifying this virus, we observe the proximal clustering of the WSEIV HA and NA to influenza B virus HA and NAs **(Fig. 1A and 1D)**. This encompasses the seasonal vaccine strains from the two influenza B virus antigenic lineages, B/Victoria/2/1987-like and B/Yamagata/16/1988-like, and the ancestral pre-divergence B/Lee/1940 virus too.^11^ The sequences of each glycoprotein were superimposed onto the publicly available structure of influenza B/Brisbane/60/2008 virus counterparts to visualize where the ~45% (HA) and ~48% (NA) identity is present **(Fig. 1B and 1E)**. As a comparative control, influenza B/Malaysia/2506/2004 (part of the B/Victoria/2/1987-like lineage) virus was selected and the HA and NA of this virus was used for the experiments detailed in this study. A pair-wise alignment of the WSEIV HA and NA with the influenza B/Malaysia/2506/2004 virus was carried out **(Fig. 1C and 1F)**. In context of the HA, there appeared to be mismatches in the residues that constitute the sialic acid interacting receptor binding site.^12,13^ This lack of conservation was indicative of a potentially altered receptor binding profile of the WSEIV HA. Additionally, the WSEIV HA also appeared to have a reduced number of putative N-linked glycosylation sites as identified by the consensus sequence N-X-(S/T). From an antigenic standpoint, the target epitope of the pan-influenza virus HA mAb, CR9114, was found to have some mismatches too.^14^ A large number of matched residues, however, appear to be in the stalk domain of the HA, consistent with previous studies showing higher levels of conservation in this region across all influenza virus HAs.^15^ Significant mismatches were observed in the region immediately upstream to the fusion peptide, largely conserved, encompassing the proteolytic cleavage site essential for activation of the HA. The comparison of the NA sequences on the other hand, demonstrated a conserved enzymatic active site, as are the regions in its immediate vicinity.

**Figure 1.**
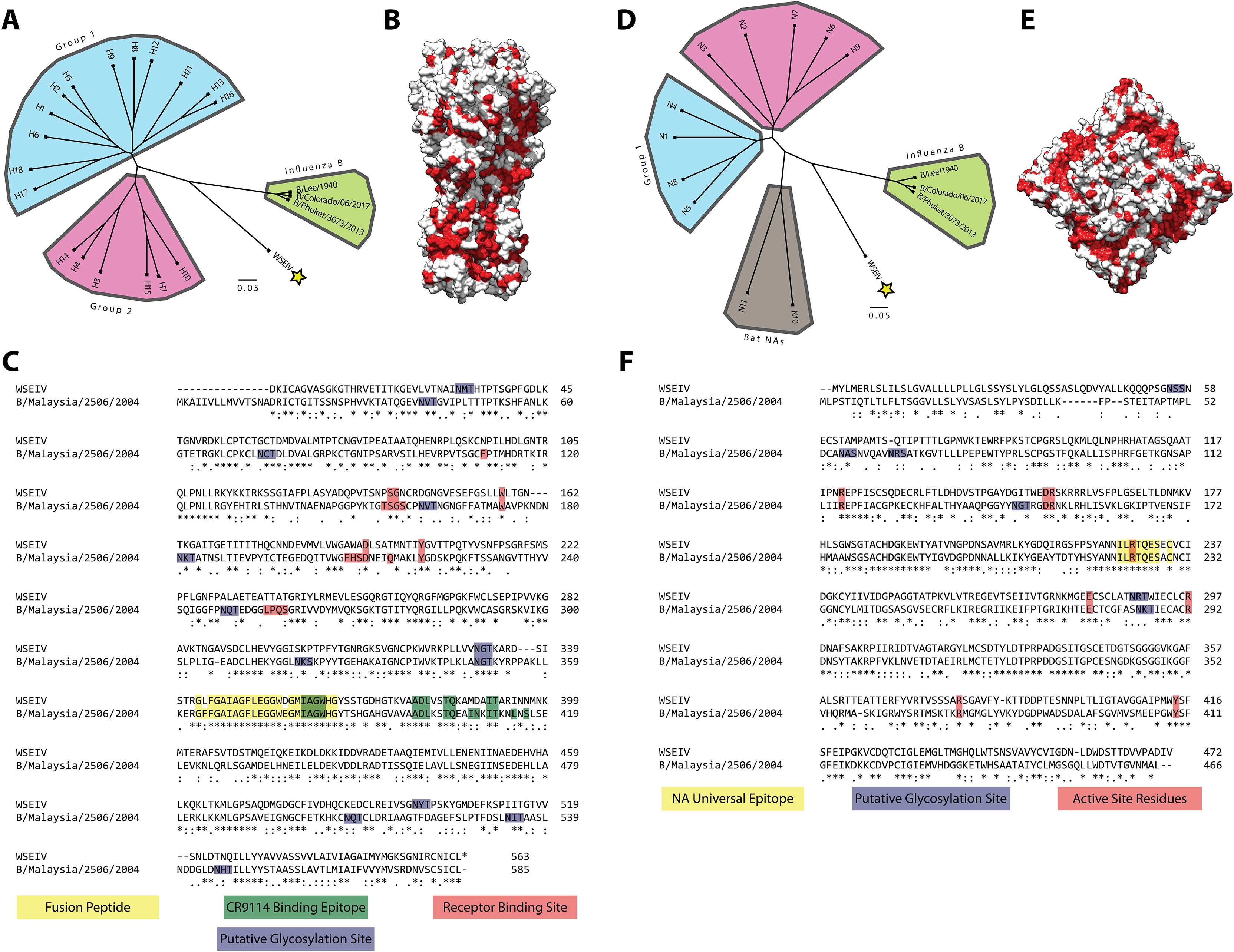
Comparative sequence analysis of the WSEIV glycoproteins. Phylogenetic trees based on the amino acid sequences of the WSEIV glycoproteins and the representative HAs **(A)** and NAs **(D)** obtained from GISAID are shown. The scale bar indicates a 5% difference in amino acid sequence. Amino acid conservation of the WSEIV HA **(B)** and WSEIV NA **(E)** relative to influenza B virus HA and NA are represented by the residues highlighted in red. The structure and sequence of Influenza B/Brisbane/60/2008 was used as a template for comparison. Pairwise alignments of the WSEIV HA and NA are displayed against influenza B/Malaysia/2506/2004 HA and NA, **(C)** and **(F)** respectively. Identical residues are indicated by asterisks. Functionally and antigenically relevant features have been annotated. Structures are based on PDB# 4FQM^48^ for HA and PDB# 4CPL^49^ for NA.

The WSEIV HA and NA were studied and characterized in the form of recombinant proteins as opposed to in a viral backbone for several reasons. The available sequences for these proteins comprised of only the coding regions from the study identifying them. Consequently, the packaging sequences in the non-coding regions essential to rescuing these glycoproteins in an influenza B virus backbone were not available. More importantly, introducing novel glycoproteins into a known human pathogen e.g. human-adapted influenza B virus backbone possesses a biosafety risk. As a result, these proteins were studied in a recombinant form to ensure to meet safety concerns. The B/Malaysia/2506/2004 and WSEIV HA and NAs were expressed using a baculovirus expression system as previously described.^16^ Since the available sequence for the WSEIV HA consisted of a truncated signal sequence, a full-length signal peptide from the B/Malaysia/2506/2004 virus HA was added instead. A C-terminal T4 trimerization domain was used to ensure the expression of the HAs in their native trimeric state. An N-terminal vasodilator stimulating phosphoprotein (VASP) tetramerization domain was used to maintain tetrameric structures of the respective neuraminidases.

To determine whether the WSEIV HA is capable of hemagglutinating erythrocytes, a conventional hemagglutination assay was carried out. Recombinant WSEIV and B/Malaysia/2506/2004 HA were incubated with 0.5% chicken and turkey erythrocytes starting at a concentration of 10μg of recombinant HA. The B/Malaysia/2506/2004 HA caused hemagglutination of both chicken and turkey RBCs while an absence of hemagglutination was observed with the WSEIV HA at comparable concentrations **(Fig. 2A)**. Although this has been observed for recent H3N2 virus isolates, we were intrigued by the lack of hemagglutination and questioned the receptor usage of this WSEIV HA.^17^ The receptor binding specificity of the hemagglutinin protein of influenza viruses is a vital parameter in addressing the potential of these viruses to cross species barriers and determining host and tissue tropism for infection.^18^ Glycan arrays are an instrumental tool in identifying the target receptor specificity for influenza virus HAs in determining whether they preferentially bind to α2,3-linked or α2,6-linked sialic acid receptors. We took advantage of this approach to further interrogate interactions between WSEIV HA and its potential binding partners. As a control, the recombinant H5 HA from A/Vietnam/1203/04 H5N1 influenza A virus, influenza B/Malaysia/2506/2004 rHA, and the WSEIV rHA were applied to a glycan array comprising of a variety of asialo, α2,3-linked, α2,6-linked and gangliosidic structures **(Fig. 2B)**. All recombinant HAs were probed with a fluorescently tagged anti-His antibody to determine the extent of binding to different glycans present on the array. In line with previously published receptor specificity profiles for avian HAs, the H5 rHA preferentially and predominantly binds to α2,3-linked sialic acids present on the array.^19^ The rHA of influenza B/Malaysia/2506/2004 shows binding to both α2,3-linked and α2,6-linked sialic acids, which has previously been possibly attributed to the presence of a Phe95 residue in influenza B virus HA as opposed to a conserved tyrosine residue at the same site in influenza A viruses.^20,21^ The WSEIV HA showed a unique binding profile on the glycan array **(Fig. 2B)**, no binding was observed to α2,3-linked or α2,6-linked sialic acids of increasing length as seen for the influenza B virus HA. Strong fluorescence intensity indicative of binding was observed on a singular spot on the array corresponding to a ganglioside oligosaccharide, GM2. This monosialylated ganglioside has a α2,3-linked sialic acid linked to the penultimate galactose residue. To validate this interaction of the WSEIV HA and GM2, bio-layer interferometry was applied to determine a dissociation constant (Kd) and thereby binding affinity **(Fig. 2C)**. To do so, Ni-nitriloacetic acid (Ni-NTA) sensors were loaded with fixed concentrations of hexahistidine-tagged rHA proteins (10 μg/ml) and dipped in 1.5x fold serial dilutions of recombinant GM2 starting at 100 μM. After ensuring that the sensors were loaded to saturation with the hexahistidine tagged HAs, the association and dissociation kinetics of the HA-GM2 interaction was studied. An A/flat-faced bat/Peru/033/10 H18 hemagglutinin was applied as a negative control given their unconventional nature to interact with MHC class II as receptors for entry as opposed to sialic acid residues.^22^ No binding was observed for the H18 rHA and as observed previously in the glycan array, B/Malaysia/2506/2004 rHA also showed no binding to GM2. The WSEIV HA on the other hand shows excellent association and dissociation profiles in a biphasic fashion with GM2 in a dose dependent manner. Analysis of this binding allowed us to determine the Kd value of this interaction at 7.36 x 10^−7^ M, in the micromolar range, with R^2^ and χ^2^ indicating good curve fits. These findings indicate and validate that the GM2 ganglioside is a target receptor for this WSEIV HA. Evidently, the WSEIV HA has a contrasting sialic acid binding profile in comparison to the influenza B virus rHA control.

**Figure 2.**
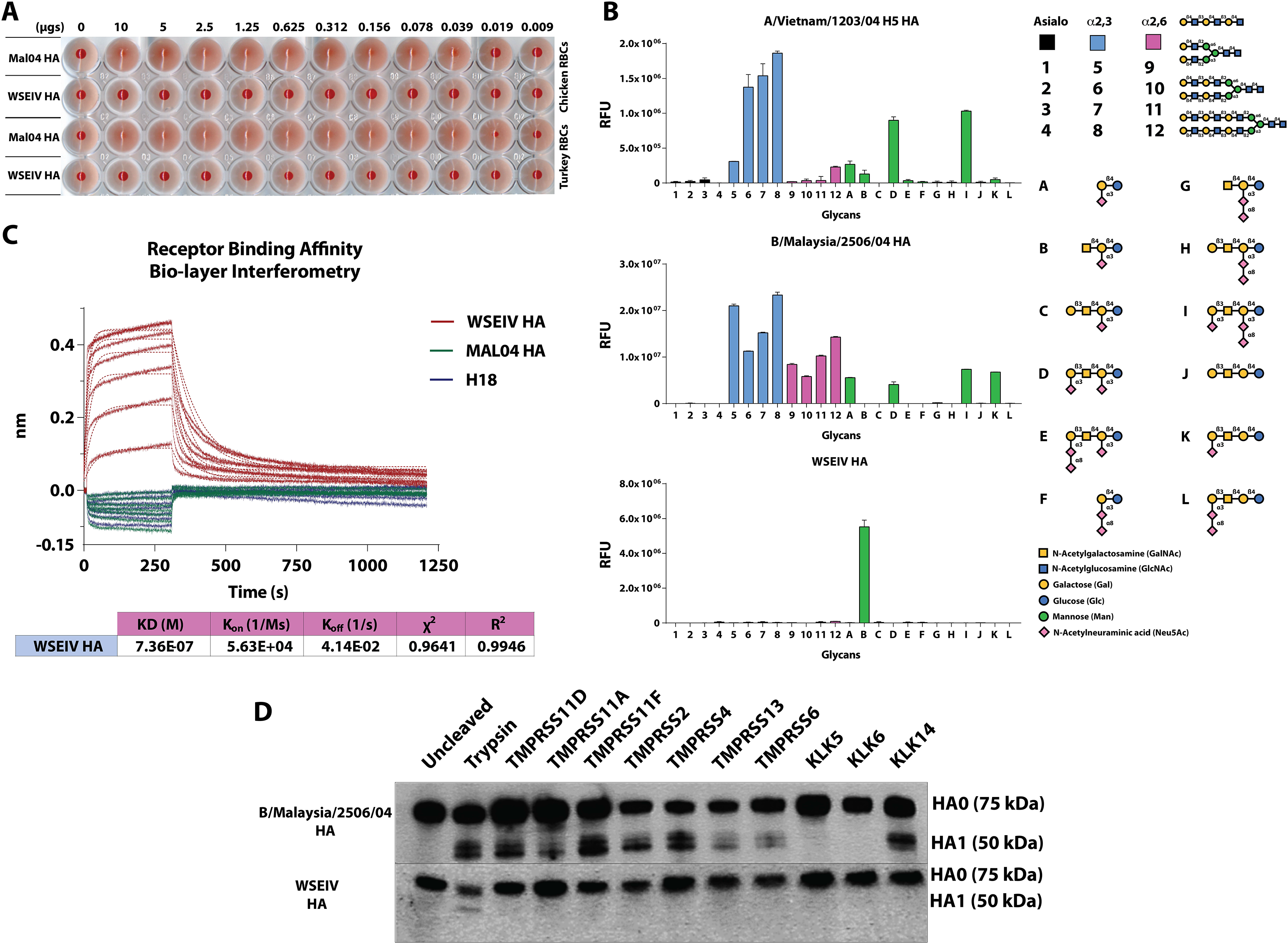
The WSEIV HA has a divergent functional profile relative to the influenza B/Malaysia/2506/2004 virus HA. **(A)** Recombinant HA protein starting at 10 μg serially diluted two-fold incubated were applied in a classical hemagglutination assay with 0.5% chicken and turkey erythrocytes. **(B)** Receptor binding specificity of the WSEIV HA was determined using a glycan microarray. A/Vietnam/1204/2004 H5 and B/Malaysia/2506/2004 HAs served as controls to compare binding profiles of the HAs. Glycans 1-4 represent asialic structures with the indicated backbones. Glycans 5-8 and 9-12 are α2,3- and α2,6-linked sialic acid glycans, with A-L comprising complex glycans and gangliosides. A simplified structure of the GM2 ganglioside, sample B on the array is shown. Error bars indicate standard deviation and technical replicates were performed. The binding was confirmed with two separate batches of recombinant WSEIV HA. **(C)** The WSEIV HA – GM2 ganglioside interaction was validated by bio-layer interferometry. B/Malaysia/2506/2004 HA and H18 HA were used as negative controls with expected absence of binding towards GM2 and sialic acids respectively. The association and dissociation kinetics curve fits of the interaction are shown and the dissociation constant (Kd) was calculated accordingly. **(D)** Western blotting was performed to determine the proteolytic cleavage profile of the WSEIV and B/Malaysia/2506/2004 HAs by human airway proteases. Cell lysate was generated following co-transfection with pCAGGS HAs and pcDNA3.1 protease expressing plasmids. HA1 was detected as a marker for proteolytic cleavage alongside uncleaved HA0. Cells transfected with pCAGGS HA either untreated or exposed to TPCK-treated trypsin served as controls for proteolytic cleavage.

Typically, following the receptor interaction of the HA with sialic acid, the HA mediates membrane fusion of the influenza virion allowing for the subsequent steps of infection and replication to occur. However, this requires the HA to be fusion competent, a state that is reliant on host cell protease-mediated activation. Cleavage by host proteases at the target site on the HA upstream of the fusion peptide splits the HA0 precursor into the HA1 and HA2 polypeptides.^23^ Recently, a study dissected the difference in preferential proteolytic cleavage of influenza A and influenza B virus HAs by human airway proteases^24^. This is crucial in addressing host adaptation of HAs and instrumental in crossing of species barriers with relative ease as observed with highly pathogenic avian influenza viruses containing a polybasic cleavage site, resulting in easier cleavage and priming of the HA by furin-like proteases.^25^ Influenza B virus HAs have been found to be cleaved by a broad range of human airway proteases belonging to the type II transmembrane serine protease and kallikrein protease families.^24^ We assessed the extent to which these proteases can cleave the WSEIV HA **(Fig. 2D)**. Human embryonal kidney (HEK) 293T cells were co-transfected with pCAGGS expression plasmids encoding for either the WSEIV or the B/Malaysia/2506/2004 HA and pcDNA expression plasmids encoding individual human airway proteases. The proteases selected here have been previously shown to cleave influenza B virus HAs.^24^ As an untreated control, HEK293T cells were transfected only with pCAGGS expression plasmids encoding the respective HAs. N-tosyl-L-phenylalanine chloromethyl ketone (TPCK)-treated trypsin was used as an additional control, with pCAGGS HA-transfected cells being incubated with exogenous trypsin briefly prior to harvesting of the cells. Cleavage of HA0 was detected through Western blotting after running the transfected cell lysate on a reducing sodium dodecyl sulfate polyacrylamide gel electrophoresis (SDS-PAGE) gel. A pool of monoclonal antibodies that are broadly cross-reactive to influenza B virus HAs and polyclonal sera raised against the WSEIV rHA in mice were used to detect cleavage of the B/Malaysia/2506/2004 HA and WSEIV HA respectively. Proteolytic cleavage of the B/Malaysia/2506/2004 HA was detected to varying extents with this panel of human airway proteases, evidenced by the presence of the HA1 bands (~50 kDA) in addition to the HA0 bands (75 kDA). Strikingly, none of the selected human airway proteases were able to cleave and activate the WSEIV HA. HA1 was detectable only in the trypsin treated sample **(Fig. 2D)**. Overall, this result supports the notion that the WSEIV HA is functionally divergent from the influenza B virus HA.

Given that the primary active site residues crucial for the sialidase activity of the NA are conserved, we tested the functional activity of the WSEIV NA. To do so, a conventional enzyme linked lectin assay (ELLA) was carried out using fetuin as a substrate **(Fig. 3A)**. Recombinant NAs were applied in 3 fold serial dilutions starting at 15 μg/ml and the extent of NA activity was determined based on extent of sialic acid cleavage, detected by binding of peanut agglutinin lectin to exposed galactose residues. The overnight incubation of the rNA with fetuin was carried out at 4 different temperatures, 4°C, 20°C, 33°C, and 37°C to determine the temperature dependence of the NA activity. These were selected to encompass the possible temperatures encountered by the spiny eels in their freshwater reservoir in the Wuhan area. Given that the enzymatic activity of the influenza B NA is understudied, an N9 rNA from the A/Anhui/1/2013 H7N9 virus was used as a comparative control. The major observation was that the WSEIV NA does indeed have neuraminidase activity **(Fig. 3A)**. This enzymatic activity is also starkly similar to that of the B/Malaysia/2506/2004 NA at the tested temperatures. Preliminary analysis suggests that the N9 rNA appears to have a higher enzymatic activity compared to both the influenza B virus and influenza B virus-like NAs. To better understand the temperature dependency, the specific activity of the NA was determined using the inverse of the half-maximum lectin binding. As expected, we observe a step-wise reduction in the activity of each NA at lower temperatures. The specific activity profile is also almost identical for the WSEIV NA and the B/Malaysia/2506/2004 NA.

**Figure 3.**
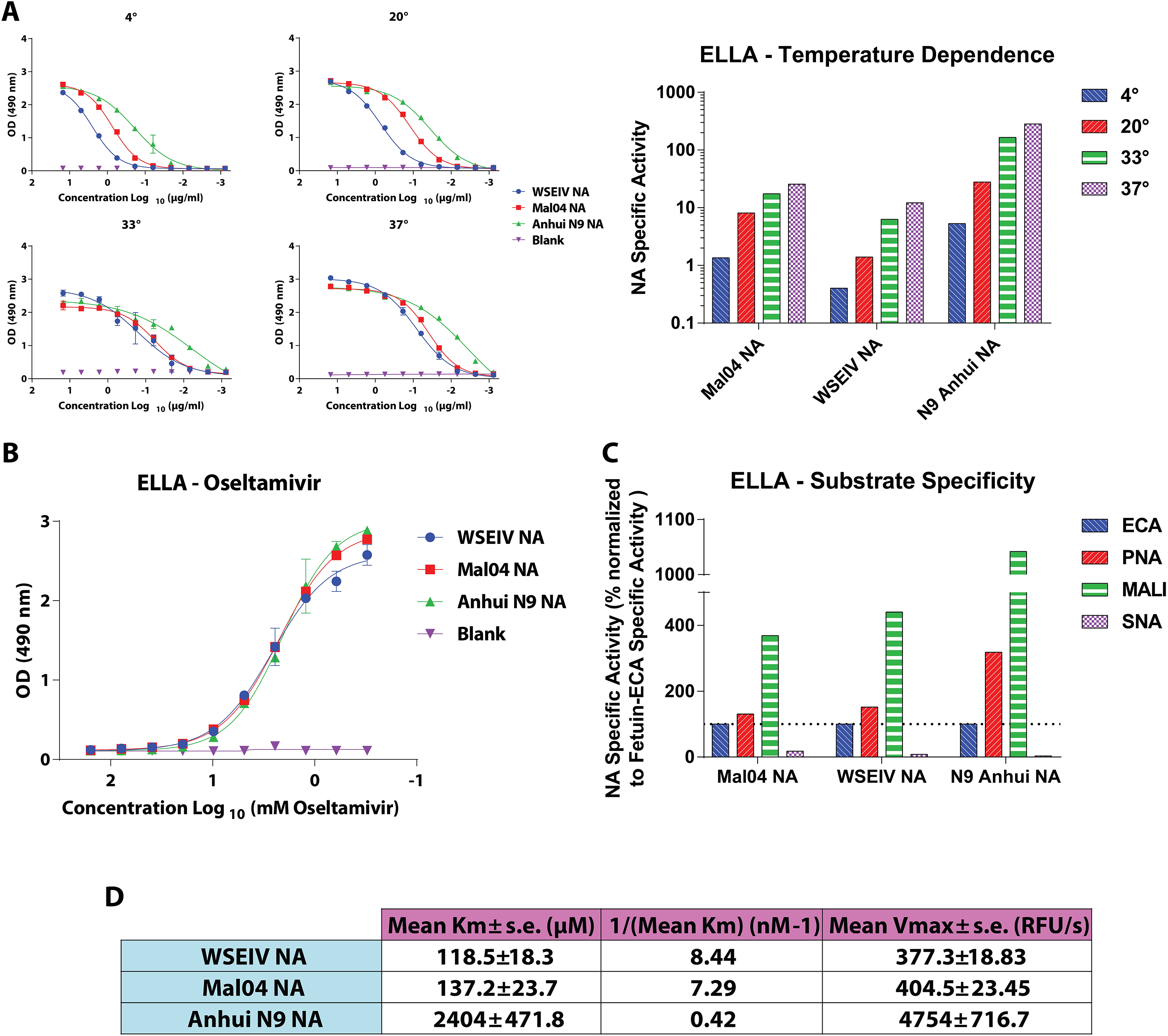
Functionally, the WSEIV NA is strikingly similar to the influenza B/Malaysia/2506/2004 virus NA. **(A)** Sialidase activity of recombinant WSEIV, influenza B/Malaysia/2506/2004, A/Anhui/1/2013 H7N9 NA proteins was evaluated in an enzyme-linked lectin assay using fetuin, at 4 temperatures; 4°C, 20°C, 33°C, 37°C. The curves indicate absorbance measured at 490 nm with error bars indicating standard deviation. Specific enzyme activity (inverse of half-maximum lectin binding) determined by these absorbance curves are shown for each individual NA at each temperature. **(B)** Susceptibility to oseltamivir was tested in a neuraminidase inhibition-based ELLA. Error bars indicate standard deviation. Absorbance at 490 nm based on lectin binding was measured with increasing concentrations of oseltamivir. **(C)** Neuraminidase substrate specificity was assessed using an ELLA using fetuin with lectins with different specificities (ECA, PNA, SNA, and MALI). Specific enzyme activity was determined as described in **(A)** and it is represented as a percentage normalized to fetuin-ECA. **(D)** Michaelis-Menten parameters, Vmax and Km, were estimated based on enzymatic activity of recombinant NAs against MUNANA substrate. Technical duplicates were performed for **(A)**, **(B)**, **(C)**, and **(D)**. The A/Anhui/1/2013 N9 NA served as a comparative control for both the WSEIV and B/Malaysia/2506/2004 NAs.

Keeping in mind the novel aspect of the WSEIV NA recently identified in a non-human host, we addressed its potential sensitivity to the NA inhibitor such as oseltamivir (Tamiflu) **(Fig. 3B)**. This was tested using a neuraminidase inhibition ELLA assay wherein fixed concentrations of rNAs were pre-incubated with oseltamivir starting at 156.28 mM and serially diluted two fold prior to being transferred onto fetuin coated plates. The extent of inhibition was estimated by increased lectin binding with lower oseltamivir concentrations. The WSEIV NA was sensitive to oseltamivir, and the dose dependency was identical for all the tested recombinant NAs **(Fig. 3B)**. These findings also align with the aforementioned observation that the active site of the WSEIV NA and the surrounding regions are well conserved to the B/Malaysia/2506/2004 NA allowing for the binding and inhibition by oseltamivir.

To follow up on the unique receptor binding profile shown by the WSEIV HA, we questioned whether the WSEIV NA has altered substrate specificity compared to the B/Malaysia/2506/2004 NA **(Fig. 3C)**. To probe this, ELLAs were carried out as previously described in a study looking at N9 neuraminidases in novel H7N9 influenza A viruses.^26^ Lectins with different binding specificities were used to determine if the NAs preferentially cleaved α2,3-linked or α2,6-linked sialic acids using fetuin as a substrate comprising of both these linkages in a 2:1 ratio.^27^ Specifically, *Erythrina crista-galli* (ECA), peanut agglutinin (PNA), *Maackia amurensis* lectin I (MALI), and *Sambucus nigra* lectin (SNA) were used. ECA and PNA both have been found to show binding to non-siaylated N- and O-linked sugars respectively, hence their binding would be higher in the presence of a NA.^28,29^ MALI and SNA preferentially bind only α2,3- and α2,6-linked sialic acids respectively, and their binding increases in the absence of NA cleavage of the target substrates^30,31^. As described earlier, specific activity of the NA was determined, and normalized to that seen with fetuin-ECA for each NA. In line with the enzymatic activity, the WSEIV and B/Malaysia/2506/2004 NA have equivalent substrate specificities, with both preferentially cleaving α2,3-linkages over α2,6-linked sialic acids **(Fig. 3C)**. Although a similar conclusion can be drawn for the A/Anhui/1/2013 N9 neuraminidase, this preferential cleavage appears to be more polarized. Subsequently, we characterized the kinetics of the enzymatic reactions of these NAs i.e. do these NAs have comparable enzyme kinetics despite having similar temperature dependent activity and substrate specificity? To investigate this, we derived the Michaelis-Menten parameters, V_max_ and K_m_, from the NA-sialic acid enzyme-substrate reaction **(Fig. 3D)**. At a fixed concentration, rNAs were incubated with the fluorogenic 4-methylumbelli-feryl N-acetyl-α-D-neuraminic acid (MUNANA) substrate and the relative fluorescence readings were taken every 90 seconds for 40 minutes as previously described^32^. The velocity of the reactions were determined for each concentration of MUNANA (starting at 1000 μM) and accordingly the V_max_ and K_m_ were calculated. The maximum enzymatic activity, V_max_, for both the WSEIV and B/Malaysia/2506/2004 NA are similar to each other, and they both display high affinity for the MUNANA substrate (indicated by the 1/K_m_). The N9 rNA shows exponentially higher maximum enzymatic activity and a markedly lower affinity for the substrate, which could possibly explain the stronger specific activity seen earlier while measuring substrate affinity.

Having characterized the functionality of the WSEIV HA and NA, we focused on surveying the extent to which broadly cross-reactive epitopes identified on influenza B virus HAs are conserved on these novel glycoproteins **(Fig. 4A and 4B)**. Probing of the WSEIV HA and NA was carried out using a panel of broadly cross-reactive human and mouse monoclonal antibodies (mAbs) previously characterized to bind to both antigenic divergent and ancestral lineages of influenza B viruses.^14,33–36^ Control enzyme linked immunosorbent assays (ELISAs) were performed for these mAbs against the influenza B/Malaysia/2506/2004 virus HA and NA and robust binding profiles for most of the tested antibodies were observed. Five antibodies showed strong binding to the WSEIV HA, namely, 1B5, 4C10, 8G3, 9B9, and 11C12 (all murine mAbs, **Fig. 4A).** With the exception of 1B5, all these mAbs bind to linear epitopes on the conserved long alpha helix in the stalk domain of the influenza B virus HA.^33^ As stated earlier, conservation between the WSEIV and B/Malaysia/2506/2004 HA (or influenza B virus HAs at large) is relatively high in this region **(Fig 1B and 1C)**. No binding of the pan-influenza virus HA human mAb, CR9114, was detectable, as anticipated from the mismatches in the binding epitope of the antibody.^14^ The binding of CR9114 to influenza B virus HA was low but detectable. Of the panel of human and mouse mAbs used to probe the WSEIV NA, only one antibody showed detectable and strong binding **(Fig. 4B)**. This human mAb, 1G01, has been characterized to have a target binding epitope in the active site of the NA with long CDR3 regions. Consequently, the breadth of the antibody encompasses influenza A and B virus NAs, showing neuraminidase inhibition activity against most of the tested influenza virus NAs.^35^ Overall, we observe an interesting profile for the WSEIV HA and NA from an antigenic standpoint. For a functionally dissimilar WSEIV HA, there are a larger number of conserved epitopes distributed in the stalk domain. Conversely, identical functionality seen for the WSEIV NA with B/Malaysia/2506/2004 NA is accompanied by an absence of this conservation outside of the enzymatic active site pocket **(Fig. 1E and 1F)**. Additionally, serum samples from humans post-seasonal influenza vaccination were used in ELISAs against the WSEIV HA and NA **(Fig 4C)**. This was done to identify basal or pre-existing cross-reactive immunity against the WSEIV HA or NA in humans as a consequence of seasonal vaccination or prior influenza virus infection. Safely assuming immunological naivety, recombinant glycoprotein from Mopeia virus, belonging to the arenavirus family was used as a target antigen in ELISAs to establish baseline reactivity in our assay. No pre-existing immunity or post-vaccination induced antibodies were detected against the WSEIV HA or NA further reinforcing the limited conservation of broadly cross-reactive target epitopes as determined by the panel of mAbs earlier.

**Figure 4.**
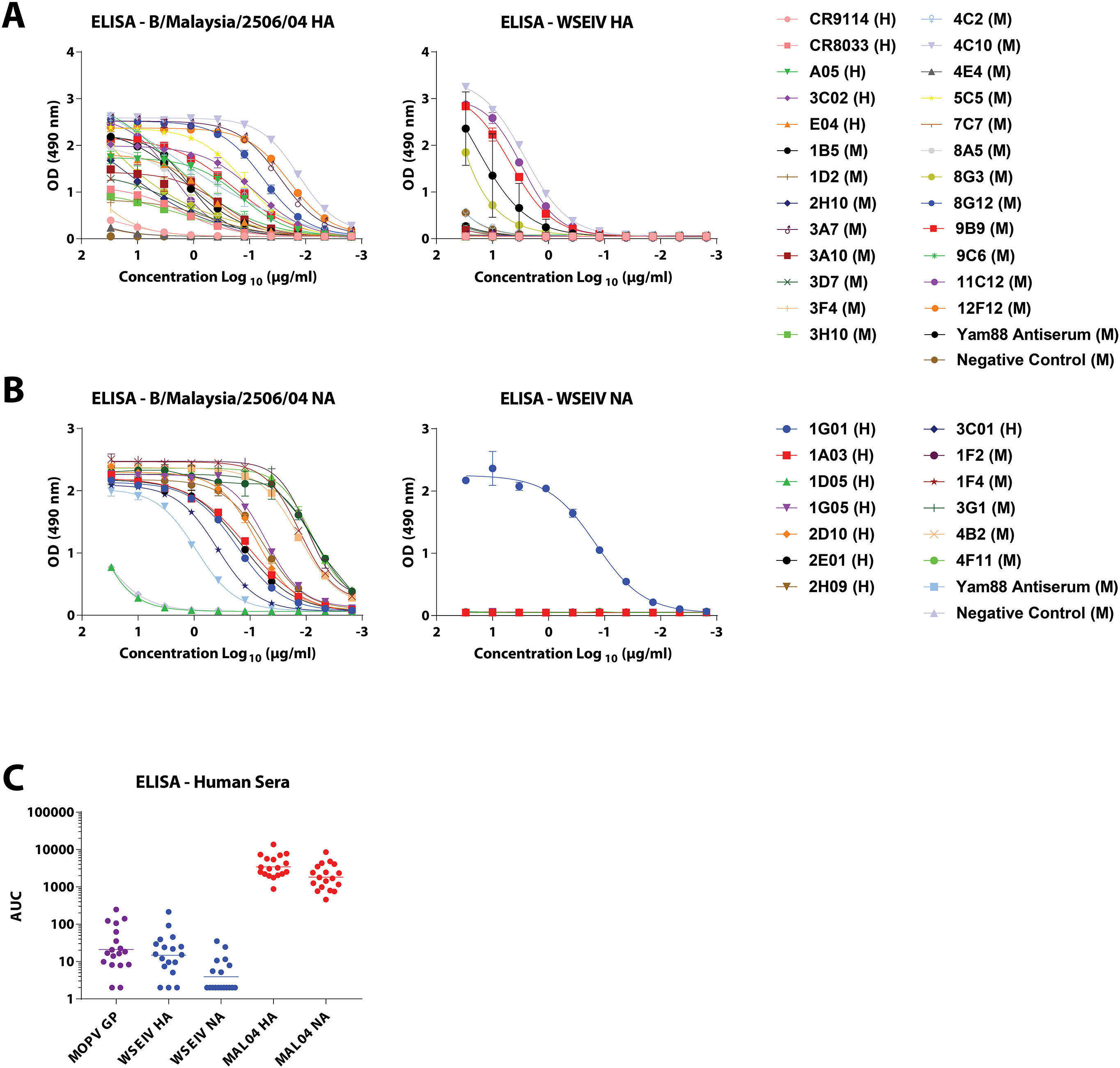
Antigenically, epitopes that are broadly conserved in influenza B virus glycoproteins occur limitedly in the WSEIV HA and NA, mostly restricted to the stalk domain of the HA and active site region of the NA. **(A)** and **(B)** Binding profiles of broadly cross-reactive anti-influenza B virus HA and NA human (H) and mouse (M) monoclonal antibodies in ELISAs are shown against recombinant B/Malaysia/2506/2004 and WSEIV HA and NAs. 4F11 (anti-influenza B virus NA) and 4C2 (anti-influenza B virus HA) mouse mAbs were used as negative controls for the ELISAs against the HAs and NAs respectively. **(C)** Presence of pre-existing immunity or induction of cross-reactive antibodies against the WSEIV HA and NA was evaluated through ELISAs using human serum samples obtained post-seasonal influenza vaccination. Mopeia virus glycoprotein was used as a negative control for baseline establishment, and area under the curve was calculated with a cutoff of average plus 3 times the standard deviation of the blank wells. The geometric mean for each group are indicated with a line.

## Discussion

Here, we have characterized the putative HA and NA of the Wuhan spiny eel influenza virus, an influenza B-like virus identified via sequence analysis in lesser spiny eels. The two studies that identify the WSEIV and a salamander influenza-like virus that also clusters close to influenza B viruses discuss the occurrence of prolonged virus-host co-divergence with several host-switching events over time.^9,37^ Largely, influenza B viruses have been discounted from being a pandemic threat due to the absence of an identified sustained non-human reservoir. Co-circulating influenza B viruses in humans have been shown to have reassortment potential within the two antigenic lineages, with B/Victoria/2/87-like viruses acquiring gene segments from B/Yamagata/16/88-like viruses.^38^ Taken together, this strengthens the unmet need for studies characterizing novel influenza B-like viruses in undersampled hosts, and subsequently understanding the functionalities of these novel viruses. Considering that vaccine approaches towards influenza viruses predominantly target the dominant surface HA, and more recently the NA glycoproteins, it is vital to have a comprehensive understanding of these proteins of influenza B-like viruses too.^39^ As well as providing this, our findings also put forth fundamental functional characterization of the influenza B virus HA and NA (sp. B/Malaysia/2506/2004) proteins. This supplements existing literature focusing on characterizing influenza B viruses which is finite in contrast to studies addressing influenza A viruses.

Along with showing limited antigenic conservation, we show that the WSEIV HA interacts with an a-series ganglioside GM2 as a target receptor. Although GM2 has been identified as an interacting partner for some reoviruses and rotaviruses, it has been shown to be not recognized by influenza A viruses, with studies demonstrating that gangliosides are entirely non-essential for influenza virus entry.^40–43^ As a therapeutic target, overaccumulation of GM2 on neuronal cells has been implicated in Tay Sachs and Sandhoff diseases with mutations rendering hexosaminidases non-functional.^44^ GM2 overexpression is also observed in a variety of human cancers and has been linked with increased tumor angiogenesis and metastatic potential.^45,46^ The WSEIV HA also appears not to bind to GM1a or GM3 as observed from the glycan array binding analysis. The receptor binding also seems to rely on the terminal GalNAc residue given that the loss of this in GM3 is accompanied by abrogation in binding. The internal location of the α 2,3-linked sialic acid residue in GM2 appears to be crucial as no binding is seen to a galactose extended GM1a ganglioside. From the perspective of gene therapy, having a viral glycoprotein that selectively targets this ganglioside could prove to be instrumental when pseudotyped into viral vectors for gene delivery. In an aquatic setting, GM2 has been found to be over expressed in gills, brain, heart, and reproductive organs of fish (zebrafish) corroborating the discovery of the WSEIV in the gills of lesser spiny eels.^47^ Having an identified receptor in fish also allows for avenues to design targeted vaccines and therapies against viruses that cause widespread economic impact in fisheries such as infectious salmon anemia isavirus. For the WSEIV NA, our findings highlight the strong conservation of enzymatic activity and kinetics, substrate specificity, and neuraminidase inhibitor sensitivity with the corresponding influenza B virus NAs. Especially the similar temperature profiles of WSEIV and influenza B virus NAs are interesting since the expectation was that the WSEIV NA would be more active at lower temperatures as found in the habitat of the lesser spiny eel. This also raises the possibility that the WSEIV is actually of mammalian or avian origin. Additional studies are needed to further determine if the virus is a *bona fide* fish virus or if it originated from other, warm-blooded animals.

Importantly, we also found very limited antigenic similarity between WSEIV and influenza B virus glycoproteins. Cross-reactivity of mAbs was limited to a small subset of antibodies and no cross-reactivity was found in human serum suggesting that we are immunologically naïve to the WSEIV glycoproteins.

In summary, the WSEIV HA and NA proteins show varying degrees of similarity to their influenza B virus counterparts. The HA displays sialic acid binding activity specifically towards GM2 and thereby differs substantially from known influenza A and B virus HAs, and the NA is indeed a sialidase with very similar functionality to influenza B virus NA. The data provided in this study contribute to our overall understanding of influenza B and influenza B-like viruses, and to understanding of the pandemic potential of the influenza B-like viruses from non-human reservoirs.

## Methods and Materials

### Cells and proteins

Human embryo kidney 293T cells were cultured in complete Dulbecco’s modified Eagle medium (DMEM; Life Technologies) constituted by DMEM supplemented with Pen-Strep antibiotics (100 U/ml penicillin, 100 μg/ml streptomycin; Gibco), 10% fetal bovine serum (FBS, HyClone), and 10 ml of 1 M 4-(2-hydroxyethyl)-1-piperazineethanesulfonic acid (HEPES, Life Technologies). Sf9 insect cells (ATCC CRL-1711) and High Five cells (BTI-TN-5B1-4 subclone; Vienna Institute of Biotechnology) were grown in Trichoplusia ni medium-formulation Hink (TNM-FH) insect medium (Gemini Bioproducts) supplemented with Pen-Strep and 10% FBS, and serum free medium (SFM)insect cell medium (HyClone) respectively.^33^

The recombinant proteins used in this study (WSEIV HA, WSEIV NA, B/Malaysia/2506/2004 HA, B/Malaysia/2506/2004 NA, A/Vietnam/1203/2004 H5 HA, A/flat-faced bat/Peru/033/2010 H18 HA, A/Anhui/1/2013 N9 NA) were expressed and purified from High Five cell culture supernatant as described in detail previously.^16^

### Phylogenetic and comparative sequence analysis

Phylogenetic trees for the HA and NA were generated as described previously.^33^ Briefly, sequences were obtained from the Global Initiative on Sharing All Influenza Data (GISAID), aligned using Clustal Omega, and the phylogenetic tree was generated using FigTree. The annotation of the tree was carried out in Adobe Illustrator. Pairwise alignment of the WSEIV and B/Malaysia/2506/2004 HA and NA was performed using Clustal Omega, following which the features of note were labeled in Adobe Illustrator. For the rendered model of glycoproteins displaying sequence conservations, the WSEIV HA and NA were aligned pairwise against the B/Brisbane/60/2008 HA and NA. The alignment was superimposed on the B/Brisbane/60/2008 HA and NA structures publicly available on PDB (HA: 4FQM^48^; NA: 4CPL^49^) using UCSF Chimera.

### SDS-PAGE and Western blotting

Recombinant proteins (10 μg) were applied to 4-20% gradient polyacrylamide gels (Bio-Rad) after heating them for 20 minutes at 95°C in 2x Laemmli buffer with 2% β-mercaptoethanol (BME). SDS-PAGE was performed at 200 volts for 35 minutes following which the gels were stained with SimplyBlue Safe Stain (Thermo Fisher) to visualize the bands alongside a color prestained protein broad range standard (New England Biolabs).

Western blotting procedures to determine the proteolytic cleavage of the HA were carried out as previously described.^24,33^ HEK293T cells were co-transfected with pCAGGS expression plasmids encoding for the corresponding HA and pcDNA3.1 plasmids encoding human airway proteases (Genscript). The Western blotting procedure was carried out with cell lysates, probing with either polyclonal sera raised against the WSEIV HA in female BALB/c mice or with a pool of anti-influenza B virus HA mAbs characterized in this reference.^33^

### Hemagglutination Assay

Recombinant HA starting at 10 μg diluted serially 2-fold was incubated with 0.5 % chicken or turkey erythrocyte suspension and incubated at 4°C for an hour. The plates were then scanned to determine the extent of hemagglutination of these erythrocytes.

### Glycan Array

Glycan array binding analysis of the rHAs was carried out as described here.^50,51^ Briefly, recombinant hexahistidine-tagged HA was precomplexed with a mouse anti-his Alexa 647 antibody (Abcam) and goat - anti-mouse Alexa 647 antibodies. This was done in 50 μL PBS-T (phosphate-buffered saline with 0.1% Tween-20) in a 4:2:1 molar ratio, incubated for 15 minutes on ice, and the applied on the array for 90 minutes in a humidified chamber. Following multiple washes with PBS-T, PBS, and deionized water the arrays were scanned to detect HA binding.

### Bio-layer Interferometry

As described previously, biolayer interferometry with an Octet Red96 instrument (ForteBio) was used to determine the dissociation constant of the HA-GM2 receptor interaction.^52^ Recombinant hexahistidine-tagged HAs at 10 μg/ml was loaded onto Ni-NTA biosensors (Fortebio) for 780 seconds to ensure saturation after a baseline step was established for 60 seconds. A second baseline was established post-loading spanning 120 seconds. Then the association (300 seconds) and dissociation (900 seconds) kinetics was recorded as the HA loaded sensors were dipped into 1.5-fold serially diluted concentrations of recombinant GM2 (Sigma Aldrich). The reaction was carried out in a 1x kinetics buffer comprising of 1x PBS, 0.01% bovine serum albumin (BSA) and 0.002% Tween 20. The dissociation constant was calculated accordingly using the suitable model for a biphasic association and dissociation profile, and global curve fit was applied to all the sensors.

### ELLAs

ELLAs were performed as described in detail previously to determine the enzymatic activity of the NAs or the oseltamivir sensitivity of the NAs.^53^ The only applied variation was that the overnight incubation was carried out at four different temperatures (4°C, 20°C, 33°C, and 37°C) to determine the temperature dependent profile of the NAs. When oseltamivir was used, the starting concentration applied was 156.28 mM with 2 fold serial dilutions, and it was pre-incubated with the recombinant NA for 1 hr shaking at 37°C after which the conventional ELLA protocol was followed. Substrate specificity characterization and specific enzyme activity determination was performed in ELLA assays identical to that described in this reference.^26^

### Michaelis Menten Kinetics

Enzyme kinetics and the Michaelis Menten parameters were determined as described previously^32^. Briefly, recombinant NAs at a fixed concentration of 10 μg/ml were incubated with 1.5 fold dilutions of the fluorogenic MUNANA substrate in MES buffer with suitable blank controls for background fluorescence. The plates were incubated at 37°C and readings for relative fluorescence unites (RFUs) were recorded at every 90 seconds for 40 minutes using a Gen5 Software in a Synergy H1 Microplate Reader (BioTek). The RFU readings were captured at excitation and emission wavelengths of 360 and 448 nM. Velocity of the reaction was determined by plotting the RFU readings against time, and the Michaelis-Menten parameters V_max_ and K_m_ were determined through non-linear regression fits of the velocity and MUNANA concentrations on Graphpad Prism 7.

### ELISAs

ELISAs were performed as previously described in detail.^33^ Information surrounding individual antibodies used in the primary staining procedure are available in the cited references.

## Acknowledgements

We would like to thank Andrew Duty and Tom Moran at the Department of Microbiology at the Icahn School of Medicine at Mount Sinai for letting us use their BLI device. We would also like to thank Edward Holmes at The University of Sidney for inspiring us to perform this study and for making the sequence information available. This work was partially funded by NIAID CEIRS contract HHSN272201400008C and NIAID grant R01 AI117287. R.P.dV is a recipient of an ERC Starting Grant from the European Commission (802780) and a Beijerinck Premium of the Royal Dutch Academy of Sciences. Synthesis and microarray analysis were funded by a grant from the Netherlands Organization for Scientific Research (NWO TOPPUNT 718.015.003) to G.-J.B.

